# Hematological and Serum Biochemical Analysis of Streptozotocin-Induced Insulin Dependent Diabetes Mellitus in Male Adult Wistar Rats

**DOI:** 10.1101/359844

**Authors:** Nima Yakhchalian, Nasim mohammadian, Kazem Hatami, Hamed Nosrati, Namdar Yousofvand

**Author notes:** Corresponding Author: Assistant Professor of Medical Physiology, Razi University of Kermanshah, Kermanshah, Iran.

## Abstract

**Background:** This investigation is concentrated on how hematological and serum biochemical markers would change in streptozotocin-induced Insulin-Dependent diabetes mellitus(IDDM) in male adult wistar rats. Hematological parameters, serum protein electrophoresis parameters and hepatic transaminases level (SGOT-SGPT) were all measured in both control group rats (N=6) and diabetic group rats (N=6) and comparison between two groups was performed.

**Material and Method:** Single dose intraperitoneal injection of 60 mg/kg dose of streptozotocin(STZ) in male adult wistar rats, induces extensive necrosis in langerhans β-cell islets, because of its cytotoxicity. Experimental diabetes mellitus can be induced completely in less than 72 hours after STZ intraperitoneal injection. Streptozotocin(STZ) was purchased from Sigma company. Diabetic and control group rats were kept separately in different metabolic cages, and their blood glucose(BG), hematological parameters, serum protein electrophoretic pattern and hepatic transaminases level were analyzed and comparison was done.

**Results:** In our investigation, Insulin-Dependent Diabetes Mellitus(IDDM) was completely induced one week after single intraperitoneal injection of 60 mg/kg BW. Diabetes mellitus induction was verified by measuring fasting plasma glucose level in blood samples of rats. Level of blood glucose, hematological parameters, serum protein electrophoretic pattern and hepatic transaminase enzymes level, were all measured. In diabetic group rats level of blood glucose (BG), hepatic transaminase enzymes (SGOT & SGPT), serum α1-globulin and β-globulin were significantly increased but in albumin, albumin/globulin ratio (A/G ratio) and serum α2-globulin a significant decrease was observed in diabetic rats in comparison with normal rats.

**Conclusion:** Extensive inflammation and tissue necrosis induced following diabetes mellitus induction in rats. Significant alterations were observed in serum protein electrophoresis fractions and hepatic transaminase enzymes level due to streptozotocin cytotoxic impacts on some tissues specifically liver.

Because of extensive β-cells necrosis and degeneration caused by streptozotocin exposure, high level of blood glucose(diabetic hyperglycemia) was observed in diabetic rats. This type of experimentally induced diabetes mellitus would highly affect hematological parameters. Insulin-Dependent Diabetes Mellitus induced by streptozotocin, can lead to anemia, neutrophilia and lymphocytosis and also has decreasing effects on red blood cell indices (HGB, MCV, MCH, MCHC) in diabetic group rats.

## 1. Introduction

One of the most imperative experimental physiology aims is to induce different types of diseases in laboratory animals and perform investigations on their various biological and pathological aspects so as to make our knowledge deeper about these diseases. In this way it would be so significant to have a holistic perspective of these diseases that has been induced in animal models in order to have a better understanding of different aspects and manifestations of these disorders. Animal laboratory medicine is crucial to have a stronger background to do research about different diseases in translational investigations. Diabetes mellitus is one of the one of the most significant and threatening complex disorders and it is needed to perform more studies on its laboratory medicine indications.

Diabetes Mellitus (DM) is an intricate metabolic disorder which can make clinical complications in the body. This disorder is identified by the pathologically elevation of blood glucose level, antioxidants decrease and abnormal metabolic pattern of lipids, carbohydrates, proteins and electrolytes(1–3). It has been approximated that nearly 366 million people are likely to become diabetic by the year 2030(4,5). The most significant diabetes mellitus complication is the extreme elevation of blood glucose level, referred as “Diabetic Hyperglycemia”, mostly induced by insulin hormone synthesis and secretion impairment, insulin function defect or a complex of these two pathological conditions(6,7). Insulin hormone is normally synthesised and secreted in full-specific, full-differentiated and full-functional pancreatic endocrine cells. Tissue-specific insulin expression is regulated at the transcriptional level, and the major regulatory elements are located in the 5′ flanking region of the insulin-encoding gene called “INS gene”(8).

Diabetic Hyperglycemia could be the result of glucose and lipid metabolic changes and also the alteration in hepatic enzymes level(1, 9). In spite of all developments in diabetology and diabetes mellitus biology and therapy, such as treatment with hypoglycemic drugs, diabetes mellitus is still a significant cause of mortality and morbidity in the world(4). Another essential diabetes mellitus complication is Reactive Oxygen Species (ROS) increase, that is the main cause of diabetes mellitus-induced hyperglycemia and can cause cell damage in many ways(10). Diabetes mellitus is widely supposed to be associated with increased oxidative stress(OS) and this could be supported by enhanced lipid peroxidase accumulation in the plasma of diabetic patients(11–16). High level of blood glucose in diabetic patients is responsible for many of the clinical manifestations such as polyphagia, polydipsia and polyuria. Diabetes Mellitus is an chronic metabolic disorder leading to critical clinical complications including chronic hyperglycemia, thirst, polyuria, visual blurriness, weight loss, lack of energy, diabetic ketoacidosis, hyperosmolar and hyperglycemic non-ketotic syndrome(17). Proteins glycation induced by chronic hyperglycemia can induce complications in kidneys, eyes, arteries and nerves. There are both non-pharmacological and pharmacological approaches in diabetes mellitus therapy. The non-pharmacological approach includes drug usage such as using oral hypoglycemic agents and insulin. Unfortunately, present conventional drugs are associated with adverse side effects like lipoatrophy, lipohyperatrophy, headache, abdominal pain, nausea, hypoglycamia, anaphylactic reaction (one of the insulin side effects) and upper respiratory tract infection(URTI) that is highly reported in the patients under treatment by metformin(18). As mentioned, none of the antidiabetic drugs can have a proper long-term glycemic control without causing any adverse side effect(19). Because of these complications in diabetes mellitus pathogenesis in the body; it needs to do more studies on this intricate metabolic disorder. Streptozotocin is a chemotherapeutic alkylating agent and a cytotoxic glucose analogue(20–23). It is the most prominent diabetogenic agent which is used to experimentally induce insulin-dependent diabetes mellitus in laboratory animals(24–28). Streptozotocin-induced diabetes mellitus is caused by extensive, severe necrosis and degeneration of the pancreatic β-cell islets(25, 29–31). This state could lead to an insulinopenia syndrome called as “Streptozotocin-induced diabetes mellitus”(21). Streptozotocin and related alkylating chemicals cytotoxic activity depends on their cellular appearance. Because of nitrosoureas high chemical affinity to lipids, streptozotocin cellular uptake through the plasma membrane is so swift and it would be accumulated in pancreatic β-cells selectively(32, 33). Insulin-producing cells which do not express GLUT2 glucose transporter on their membrane would be streptozotocin resistant(34, 35). Streptozotocin also damages other tissues expressing GLUT2 glucose transporter, specifically liver and kidney(36, 37). Streptozotocin cytotoxicity is because of its ability to have DNA alkylating and methylating activity(38, 39) and also protein glycosylation induced by this chemical agent(40). After DNA damage induced by streptozotocin cellular uptake, DNA repair mechanisms will be activated in response to DNA damages; one of the results is poly (ADP-ribose) polymerase overstimulation which can abate cellular NAD+ and stores of ATP(41–43). Cellular energy stores depletion could lead to extensive necrosis and damage of β-cells islets(44, 45). Although probably streptozotocin-induced DNA methylation and alkylation are the main responsible reasons for β-cells necrosis, it is likely that after streptozotocin exposure, β-cells functional incompetency could be developed by protein methylation. Streptozotocin was injected intraperitoneally in the dose of 60 mg/kg body weight to induce diabetes mellitus in rats. Diabetes mellitus was completely induced and subsequently, diabetic hyperglycemia was observed(46,47). The present research is focused on the effects of intraperitoneal injection of 60 mg/kg body weight streptozotocin on hematological parameters, level of blood glucose and serum protein electrophoretic pattern and hepatic transaminase levels in diabetic group rats in comparison to control group rats. In the following parts of this article we will have the literature review in second part, materials and methods in the third part and then we shall have results in the next part, and the last part would be concentrated on discussion and conclusion.

## 2. Literature Review

### 2.1 Hematology

Several studies have documented hematological alterations in streptozotocin-induced insulin dependent diabetes mellitus such as Akpan and Ekaidem research in 2015 which was concentrated on immunological and hematological alterations often occur in diabetes mellitus in result of oxidative stress induced by diabetes mellitus. Diabetes mellitus was induced by one intraperitoneal injection of 60 mg/kg BW of streptozotocin in rats. The results of this investigation showed that the diabetic control had significantly higher level of WBC count than the normal control. Red blood cell(RBC), Hemoglobin(HGB) and Packed cell colume(PCV) were all significantly reduced in comparison to normal control as well as red blood cell indices including MCV, MCH and MCHC. But platelet count(PLT) of the diabetic control was significantly increased in comparison to the control group. At the end of experimental period, the glucose level of blood received from the caudal vein of rats also showed a significant increase due to diabetic hyperglycemia(48).

A study carried out by Keskin and et al. in 2016 to estimate the beneficial and preventive effects of quercetin on some hematological parameters in streptozotocin-induced diabetic rats. Streptozotocin was injected at a single dose of 60 mg/kg (i.p) for diabetes mellitus induction. After diabetes mellitus induction, leukocyte count(WBC), erythrocyte count(RBC), hemoglobin(HGB), hematocrit(HCT), platelet count(PLT), mean corpuscular volume (MCV), mean corpuscular hemoglobin(MCH), mean corpuscular hemoglobin concentration(MCHC) and differential leukocyte count were examined in blood samples. Red blood cell(RBC) and platelet count(PLT), Hemoglobin(Hb) and hematocrit(HCT) levels in diabetic rats significantly decreased in comparison to control ones. MCV, MCH, MCHC levels did not show any significant alteration. Leukocytes count was increased in diabetic rats; the Neutrophil count was also increased in diabetic group rats(49).

Another study was Peelman and et al. research in 2004. The effective mechanism for leukocytosis in obesity, diabetes mellitus and atherosclerosis has been reported mostly unknown in that research. Recent evidence suggests that leptin and leptin receptor are parts of a pathway that stimulates hemopoiesis. This hemopoiesis stimulating pathway could be involved and lead to elevation of white blood cell count(50).

A research by Pertynska-Marczewska and et al. in 2004 showed that advanced glycation end products(AGEPs) and pro-inflammatory cytokines could be the reason for polymorphonuclears and mononuclears activation and elevation in peripheral blood. Some investigations are offering different reasons for leukocytosis occur in diabetes mellitus(51). For example in 2004 Shurtz-Swirski and et al. documented that probable mechanism of raised WBC count is oxidative stress that occur in diabetes mellitus(52)

### 2.2 Serum Clinical Biochemistry

Several studies have been carried out to evaluate serum biochemical modifications occur in diabetes mellitus such as Zafar and et al. investigation in 2009 which was performed to evaluate the impacts of Diabetes Mellitus on liver morphology, architecture and function. The hepatic effects of diabetes mellitus were evaluated in vivo using streptozotocin (STZ)-induced diabetic rats as an experimental model. Diabetes mellitus induced by a single dose of STZ (45 mg/kg, b.w.) given intraperitoneally in sodium citrate buffer at pH 4.5. Histopathological analysis of liver in diabetic rats showed accumulation of lipid droplets, inflammatory cells infilteration, increased fibrous content, dilation of portal vessels, congestion of these vessels and proliferation of bile ducts epithelial cells. Increased levels of aspartate aminotransferase (AST), alanine aminotransferase (ALT) and alkaline phosphatase(ALP), were also observed in the diabetic rats. Serum plasma glucose was also significantly elevated(46).

A comprehensive research by Zafar and et al. in 2010 revealed a plasma glucose level significant increase in diabetic rats in comparison to control group rats. Transaminase enzymes (AST and ALT) level were also increased(53).

Another research by Ahmed and et al. in 2012 on Streptozotocin-Induced Diabetes Mellitus to investigate the effects of hesperidin and naringin on serum glucose, blood glycosylated hemoglobin (HbA1C) and serum insulin levels in high fat fed (HFD)/streptozotocin (STZ)-induced type 2 diabetic rats. Also the effects on serum lipid profile, adiponectin and resistin levels, cardiac functional biomarkers and liver and muscle glycogen contents were evaluated. The significant increase of blood glucose level was also observed in diabetic rats. They reported a significant elevation in AST, ALT, LDH and CK-MB in diabetic rats as well(54).

In Kim et al. (2014) study diabetes mellitus was induced by single injection of streptozotocin (STZ, 200mg/kg, i.p.) in mice. After diabetes mellitus induction, plasma glucose, SGOT, SGPT, LDH and ALP were significantly increased in diabetic group mice(55).

Salih et al. in 2014 designed a research to evaluate the biochemical changes of liver function following STZ-induced Diabetes Mellitus in mice. They induced diabetes mellitus by a single injection of STZ (150mg/kg BW i.p.) then serum samples were collected after two, four and six weeks of injection. Significant increase in blood glucose level, the liver weight/body weight ratio, levels of ALP, AST, ALT, bilirubin and cholesterol were observed. Increase in the activity of ACP was insignificant. Serum LDH was increased significantly by the second and fourth week but decreased at the sixth week. The study concluded that STZ-induced Diabetes Mellitus affects the biochemical function of liver and causes liver enzymes disturbances (56).

Single dose intraperitoneal injection of 60 mg/kg streptozotocin in male adult wistar rats causes extensive necrosis in langerhans β-cell islets and in less than 72 hours can induce experimental diabetes mellitus. Streptozotocin was purchased from Sigma company. The diabetic and normal rats were kept in metabolic cages separately, and their hematological parameters, serum protein electrophoretic pattern and level of AST and ALT were measured in all animals after the experimental period, and the comparison between these groups was done. Intraperitoneal injection was done in non per oral state. Streptozotocin was dissolved in 10 mM sodium citrate buffer before giving intraperitoneal injection. Twenty male adult wistar rats(200-250 gr) were used for this investigation, housed in controlled temperature room (20-22 °C) with a 12 h light/dark period. All animals were kept according to criteria outlined in the guide for the care and use of laboratory animals prepared by national academy of sciences and published by national institute of health. All procedures were performed in sterilised condition. These twenty male adult wistar rats were divided into two seperate groups including control (n=6) and diabetic group (n=6). Streptozotocin was dissolved in citrate buffer (15 mg/ml) 4.5 PH solution. Control rats (n=6) were given only physiological saline (50 mg/kg bw i.p.) and streptozotocin-induced diabetic rats(n=6) were exposed to 60 mg/kg body weight by intraperitoneal administeration. Blood samples were prepared for hematological analysis by collecting blood directly from the heart. Collected blood was allowed to clot for about 30 minutes at room temperature(20-22 °C) and serum was obtained from fresh blood of rats to measure biochemical parameters. The hematological parameters, serum protein electrophoresis parameters and hepatic transaminases level were measured after seven days of diabetes mellitus induction. In all of the groups, the blood samples were centrifuged at 3000 rpm for 15 min, plasma and buffy coat were separated then samples were obtained and analyzed.

## 3.1 Hematological Assay

Blood samples were collected directly from the heart. The complete blood count (CBC) was performed on an automated hematology analyzer using well-mixed whole blood with K2-EDTA to prevent blood clotting. CBC/diff analysis was performed by hematology diagnostics division. Total white blood cell count (WBC) was estimated according to the visual method of Dacie and Lewis (1991)(58). The percentage of packed cell volume (PCV) was determined according to the hematocrit method, while the blood hemoglobin (Hb) concentration in all blood samples was measured according to the cyanomethemoglobin method using Drabkin’s reagent (Alexander and Grifiths,1993)(59). Mean cell volume (MCV), mean corpuscular hemoglobin (MCH) and mean corpuscular hemoglobin concentration (MCHC) were calculated as outlined in Dacie and Lewis (1991). Differentiated white blood cell counts were estimated using the method of Osim et al.(60).

## 3.2 Biochemical Assay

### 3.2.1 Blood Glucose Level

At the end of the experimental period, rats blood glucose concentration were checked by caudal vein blood sampling and using Bionime glucometer.

### 3.2.2 Serum Protein Electrophoresis(SPEP)

Serum protein electrophoresis (SPEP) was performed on cellulose acetate strips by using an already made buffer (pH 8.6). The cellulose acetate layers have initially been soaked in the buffer solution, and more amounts of the buffer was removed by placing them in between two Whatman no-1 filter papers. Then, the strips were put on the central compartment of the electrophoresis chamber. Two filter paper strips were placed on both sides of the cellulose acetate strip to connect them with the two buffer containing chambers on both sides of the electrophoresis chamber. Then, ten microliters of the serum samples were loaded on the cellulose acetate strip at the sources of the origin. Then, the electrophoresis chamber was connected to the power pack, and it was subjected to electrophoresis. After one hour, the strips were removed, and they were stained by using Ponceau S. after distaining them by using the reagent. Finally, the separated protein fractions could be observed.

### 3.2.3 Hepatic Transaminase (AST-ALT)

The activities of serum aspartate aminotransferase (AST) and alanine aminotransferase (ALT) were estimated using standard methods on Cobas MIRA_ (Roche Diagnostics, Basel, Switzerland) automated analyser.

## 4. Results

Following single intraperitoneal injection of 60 mg/kg body weight streptozotocin, diabetes mellitus was induced completely after one week, then blood glucose level, hematological parameters, serum protein electrophoresis parameters and hepatic transaminases were measured to perform a comparison between normal control and diabetic control rats. All data are scheduled in the following tables.

**Table no 1:** Showing Hematological Analysis of Control and Diabetic Rats

| Group | WBC | HGB | MCV | MCHC | MPV | PLT | Neutrophil | Lymphocyte | MCH | HCT % |
| --- | --- | --- | --- | --- | --- | --- | --- | --- | --- | --- |
| Control | 2560.00<br>±<br>620.08 | 15.88<br>±0.49 | 56.51<br>± 0.584 | 33.03<br>± 0.29 | 6.45<br>± 0.15 | 183833.33<br>±<br>114979.24 | 530.00<br>±124.01 | 1987.50<br>± 465.06 | 18.73<br>±0.28 | 37.23<br>±2.41 |
| Diabetic | 5966.67<br>±<br>1249.17 | 11.25<br>±0.84 | 50.33<br>± 1.09 | 27.11<br>± 1.88 | 6.50<br>± 0.22 | 366000.00<br>±<br>95645.23 | 1193.33<br>±249.836 | 4460.00<br>± 936.443 | 16.33<br>±0.51 | 42.65<br>±3.69 |
| Change percentage from control to diabetic rats | +133.04 | -29.15 | -10.93 | -17.92 | +0.77 | +99.09 | +125.15 | +124.40 | -12.81 | +14.55 |

**Table no 2:** Showing Biochemical Analysis of Control and Diabetic Rats

| Group | Glucose(mg/dL) | AST(IU/L) | ALT(IU/L) | Albumin | α-1 Globulin | α-2 Globulin | β-Globulin | A/G ratio |
| --- | --- | --- | --- | --- | --- | --- | --- | --- |
| Control | 105.33±5.77 | 81.75±9.37 | 60.00±4.88 | 71.25±1.09 | 7.00±0.16 | 14.85±1.53 | 10.45± 0.50 | 2.38±0.17 |
| Diabetic | 374.33±34.81 | 137.167±19.22 | 120.66±11.54 | 51.61±3.27 | 9.88±1.10 | 9.70±0.92 | 18.30±2.67 | 1.13±0.19 |
| Change percentage from control to diabetic rats | +255.38 | +67.77 | +101.1 | -27.56 | +41.14 | -34.68 | +75.11 | -52.52 |

All data are expressed as Mean±SEM and p < 0.05 considered significant difference when diabetic group rats compared with control group rats.

### 4.1 Hematology

Hematological alterations in diabetic group rats were observed. Streptozotocin-induced insulin-dependent Diabetes Mellitus can lead to HGB decrease (anemia), neutrophilia (significant elevation in Absolute Neutrophils Count) and lymphocytosis (significant increase in Absolute Lymphocyte Count). The total number of leukocytes (leukocytosis) was observed in diabetic rats in comparison to normal control group rats. RBC indices (MCV, MCH, MCHC) were also decreased significantly in diabetic rats. There was no significant difference (P>0.05) observed in Platelet count (PLT) and mean platelet volume (MPV) between two groups.

### 4.2 Biochemistry

Biochemical analysis was done on serum samples in both diabetic and control group rats. Level of fasting plasma glucose (FPG) was significantly elevated in diabetic rats in comparison to normal rats. Serum protein electrophoretic pattern (SPEP) was significantly changed in diabetic group rats. Significant alteration in serum proteins observed in diabetic rats including significant α-1 globulin and β-globulin fractions elevation (p<0.05), and α-2 globulin and albumin fractions significant decrease (p<0.05). Albumin/Globulin ratio (A/G ratio) was also decreased significantly (p<0.05) in diabetic rats.

## 5. Discussion

Diabetes mellitus (DM) is recognized as a complicated metabolic disorder described by lack of blood glucose concentration homeostasis and abnormal metabolic pattern of carbohydrates and lipids(61). Persistent hyperglycemia during diabetes mellitus can induce many chemical alterations such as nonenzymatic glycosylation of proteins in the body (62) or generating Reactive Oxygen Species(ROS)(10). These chemical mechanisms in diabetes mellitus are responsible for most of the clinical complications observed in diabetic patients. Alteration in the activity of several glycolytic and gluconeogenic enzymes of the liver have also been documented in diabetic state(1). Streptozotocin-induced diabetes mellitus provides chronic oxidative stress along with the resulting hyperglycemia(63). Streptozotocin is a cytotoxic alkylating agent which induces swift and irreversible necrosis in pancreatic β-cell islets(64).

Whereas a single diabetogenic dose of streptozotocin has been used to induce complete β-cells degeneration in most species within 24 h, multiple sub-diabetogenic doses of streptozotocin partially damage β-cell islets, moreover induced inflammatory process could lead to macrophage and subsequent infiltration of lymphocytes which would be followed by the insulin deficiency onset(65).

Earlier analyses have used different doses of streptozotocin to induce diabetes mellitus in laboratory animals. The single diabetogenic dose of streptozotocin has been documented in different studies from 50 mg/kg body weight(46) up to higher doses such as 70 mg/kg(66) or intraperitoneal injection of 150 mg/kg body weight(56). These doses could be used to induce diabetes mellitus completely.

One week after intraperitoneal injection of streptozotocin, diabetes mellitus was induced completely in rats. Fasting plasma glucose (FPG) showed a significant increase in diabetic group rats(p<0.05), so this investigation confirmed previous studies(67).

Proteins glycation induced by diabetic hyperglycemia can have modificational impacts on erythrocytic membrane proteins and hemoglobulin(Hb) as well(68). Proteins dysfunction in diabetes mellitus could be induced by these biochemical modifications and probably are responsible for long-term diabetes mellitus clinical complications. Lipid peroxidation increase, is one of the most significant findings in diabetes mellitus, which is a marker of elevated oxidative stress in diabetes mellitus(69). Increase of lipid peroxides induced by elevated lipid peroxidation, and glycosylation of cell membrane proteins may lead to red blood cells membrane damage, hemolysis and anemia. Free radicals-induced lipid peroxidation of the membrane could increase the rigidity of plasma membrane and decrease deformability of cells, reduce erythrocytic survival and fluidity of lipids(70).

The relation between anemia and chronic disorders has been well-documented and confirmed in different researches such as Weiss et al. in 2005. In this research, the RBC membrane lipid peroxide was not measured however erythrocytic indices including MCV, MCH, MCHC and HGB were measured(71). The MCH, MCHC and HGB values in diabetic group rats showed significant decrease in accordance with prior hematological studies reporting anemia as a pathophysiological complication of diabetes mellitus(72). This reduction is probably because of blood osmoregulation defect and abnormal synthesis of hemoglobin (HGB)(73). HCT was not significantly changed in diabetic rats in comparison to normal rats (p>0.05). Mean platelet volume (MPV) and platelet count (PLT) were also measured, and both hematologic markers were not significantly changed in diabetic rats in comparison to normal rats(p>0.05).

Serum ferritin is a biomarker that can reflect total body iron stores; moreover it is a sensitive biomarker of inflammatory stress in the body(74). Serum ferritin increase has been reported in diabetes mellitus that could reveal inflammatory component of this complex metabolic disorder(75), and this inflammatory component can affect lots of biomarkers in the body such as hematological parameters and serum biochemical factors.

In the peripheral blood analysis, leukocytosis (absolute increase in the white blood cells count) in both of the polymorphonuclears and mononuclears count was observed in diabetic group rats. It could not be only the result of oxidative stress in diabetes mellitus(52). It also could be induced by glycation end products. Glucose interaction with proteins amino groups, produces derivatives called “advanced glycation end products (AGE)”, it has been demonstrated that these glycation end derivatives could enhance some angiogenic and inflammatory cytokines expression including VEGF, TNF-α and IL-8, and this process could be the reason for leukocytes activation and increase(51). In other words, Transformation growth factor (TGF)-β1(76), Superoxide(77), interleukin-1β and other chemokines(78) can activate leukocytes to take part in diabetes mellitus pathogenesis. The exact mechanism of leukocytosis in diabetes mellitus is still unknown, but it is suggested that leptin and leptin-receptors are parts of a hemopoiesis stimulating process(50). Interestingly, the high structural similarity between leptin-receptors and class-1 cytokine receptors has been shown(79) Therefore, there is probably a similar pathway for IL-6 and leptin playing their role and this can probably explain why leukocytes are enhanced in diabetes mellitus(80).

Earlier studies on hematological analysis in streptozotocin-induced diabetes mellitus have revealed moderate neutrophilic leukocytosis in diabetic rats(81). Our investigation is also in accordance with these studies.

Serum biochemical assay including serum protein electrophoresis analysis and hepatic transaminase enzymes level evaluation were also done on serum samples of both control and diabetic group rats. Electrophoresis of serum proteins showed the level of different bands of serum protein fractions including albumin, α-1 globulin, α-2 globulin, β-globulin and γ-globulin in the blood. Serum protein electrophoretic pattern was changed in diabetic group rats. Alteration pattern in serum proteins observed in diabetic rats, including significant α-1 globulin and β-globulin fractions elevation (p<0.05) and α-2 globulin and albumin fractions significant decrease (p<0.05). Albumin to globulin ratio(A/G ratio) was also decreased significantly (p<0.05) in diabetic rats.

It has been shown that hepatobiliary diseases such as necrosis, inflammation or non-alcoholic hepatic steatosis could be induced by diabetes mellitus(82, 83) and the mortality rate caused by advanced-stage hepatic diseases in diabetic patients is higher than cardiovascular causes(84), therefore hepatic injuries can develop in diabetes mellitus and these hepatocellular injuries probably play striking role in serum protein electrophoretic pattern alteration in diabetes mellitus.

Some mechanisms such as oxidative stress, chronic hyperglycemia, chronic inflammation, lipotoxicity and many other mechanisms can induce pancreatic β-cells dysfunction in diabetes mellitus(85, 86). An investigation has demonstrated that oxidative stress and hepatocellular fat accumulation (hepatic steatosis) can play a significant role in hepatic disorders caused by diabetes mellitus(87). It has also been investigated that endoplasmic reticulum stress(ERS) is another important factor in pancreatic β-cells dysfunction in diabetes mellitus(88). It has also been investigated that endoplasmic reticulum stress (ERS) is another critical factor in pancreatic β-cells dysfunction in diabetes mellitus(88). Pro-inflammatory cytokines such as TNF-α and interleukin-1β (IL-1β) can activate signalling cascade of cell death including ASK-1 and p38 MAPK in the diabetic liver and this could lead to hepatocellular damage in diabetes mellitus(89). These hepatic damages induced by diabetes mellitus could lead to many alterations in serum proteins concentration especially serum protein electrophoretic pattern.

Inflammatory component of diabetes mellitus has been investigated and proved in many studies. The significant change in the cocentration of acute phase reactants (APR) in serum protein electrophoresis profile was observed in diabetic group rats. APRs are essential glycoproteins involved in inflammatory responses in the body(90). α-1 antitrypsin, α-1 acid glycoprotein, β-globulins, plasma fibrinogen, C-reactive protein (CRP) and many others are known as “positive acute phase reactants” which will increase in inflammatory response and albumin, pre-albumin and transferrin are known as negative acute phase reactants which show decrease in inflammatory responses in the body. The response to inflammation by acute phase reactants is called as “Acute Phase Response”. Synthesis of these acute phase reactants(APRs) which mostly done by liver can be enhanced in response to specific cytokines secreted in inflammatory processes such as IL-1 and TNF-α(91, 92) and as mentioned, these cytokines(TNF-α, IL-1β) are elevated in diabetes mellitus because of its inflammatory nature(51, 89). Serum albumin as explained is a negative acute phase reactant, therefore decrease of serum albumin in diabetic rats could be induced by inflammatory processes involved in the pathogenesis of streptozotocin-induced diabetes mellitus as explained(75). Hepatic necrosis and inflammation induced by diabetes mellitus pathogenesis is probably another reason for albumin level decrease in the blood(87, 89) because hepatocytes are the only producing units of albumin protein in the body and hepatic injuries can decrease albumin level in blood because of hepatocytic damages. Inflammatory mechanisms involved in diabetes mellitus also can affect globulins level in blood as discussed. α-1 globulin and β- globulin elevation was observed due to inflammatory response or because of streptozotocin-induced diabetes mellitus resulting extensive necrosis(93–95).

α-2 globulin and albumin/globulin ratio(A/G ratio) markers were also analysed. α-2 globulin and albumin/globulin ratio (A/G ratio) were decreased significantly (p<0.05). α-2 globulin decrease could be the result of hemolysis due to elevated lipid peroxides and red blood cells membrane damage(70) or hepatic dysfunction induced by diabetes mellitus, because the liver is responsible for haptoglobulin synthesis that is a significant component of α-2 globulin fraction in serum protein electrophoresis(93–95).

Hepatic transaminases including AST and ALT were also measured in blood samples. Both enzymes were increased significantly in diabetic rats blood(p<0.05). This could be the result of hepatic necrosis development in diabetes mellitus and hepatotoxicity.

Both hepatic lesions or even hemolysis could induce AST elevation because it could be also found in red blood cells; therefore, the hemolytic process could be a reason for its increase in blood samples, but ALT enhancement is more liver-specific and shows hepatocellular extensive damage in diabetic group rats.

Many studies have reported that single diabetogenic dose of streptozotocin showed an increase in glucose, ALT and AST levels(46, 53–56).

The present study results confirmed the previous studies and showed that single dose of streptozotocin causes a significant increase in glucose level in diabetic rats in comparison with control group rats and also showed an increase in ALT and AST.

## 6. Acknowledgments

We are extremely grateful to our friends and advocators, PANTEA BAZEGHI, DANIAL KHOSHKHO, MASOUD NOSRATI and Madar Clinical Pathology Lab in Kermanshah. We are also wholeheartly appreciated to everyone who helped us to carry out this investigation.

## 7. Conclusion

The present study was carried out to investigate hematological and serum clinical chemistry alterations in experimental animal model of diabetes mellitus induced by single intraperitoneal injection of 60 mg/kg body weight streptozotocin. At the end of the experimental period, the significant elevation in the level of blood glucose(BG), hepatic transaminases including Aspartate aminotransferase (AST) and Alanine aminotransferase (ALT) was seen. In serum protein electrophoresis analysis, serum α-1 globulin and β-globulin fractions showed significant increase but serum α-2 globulin fraction, albumin and albumin/globulin ratio showed significant decrease. Also, many hematological changes were observed in diabetic group rats. RBC indices including MCH, MCV, MCHC and HBG were significantly decreased in diabetic rats, but there was no significant change in HCT, MPV and PLT in diabetic rats in comparison with control group. Total white blood cells increase (leukocytosis) was also observed in diabetic group rats.

## References

1. Anderson J, Stowring L. Glycolytic and gluconeogenic enzyme activities in renal cortex of diabetic rats. American Journal of Physiology. 1973;244:930–6.

2. Hink U, Li H, Mollnau H, Oelze M, Matheis E, Hartmann M, et al. Mechanisms Underlying Endothelial Dysfunction in Diabetes Mellitus. Circulation Research. 2001;88(2).

3. Jay D, Hitomi, Griendling K. Oxidative stress and diabetic cardiovascular complications. Free Radical Biology and Medicine. 2006;40(3):183–92.

4. Nammi S, Boini MK, Lodagala SD, Behara RBS. The juice of fresh leaves of Catharanthus roseus Linn reduces blood glucose in normal and alloxan diabetic rats. BMC Complement Altern Med. 2003;3:1–4.

5. Wild S, Roglic G, Green A, Sicree R. Global prevalence of diabetes: estimates for the year 2000 and projections for 2030. American Diabetes Association. 2004;27(5).

6. Mohammed A, Tanko Y, Okasha M, Magaji R, Yaro A. Effects of aqueous leaves extract of Ocimum gratissimum on blood glucose levels of streptozotocin-induced diabetic Wistar rats. Afr J Biotechnol. 2007;6(18):2087–90.

7. Nakhaee A, Bokaeian M, Saravani M, Farhangi A, Akbarzadeh A. Attenuation of oxidative stress in streptozotocin-induced diabetic rats by eucalyptus globules. Indian J Clin Biochem. 2009;24:419–25.

8. Ohneda K, Ee H, German M. Regulation of insulin gene transcription. Semin. Cell Dev Biol. 2000;11:227–33.

9. Jenson T, Deckert T. Abnormalities in plasma concentration of lipoprotein and fibrinogen in type 1 (insulin dependent) diabetic patients with increased urinary albumin excretion. Diabetologia. 1998;31:142–6.

10. Har H, Ja AS. Endothelial dysfunction in diabetes mellitus. Vascular health and risk management. 2007;3(6):853–76.

11. Hunt J, Dean R, Wolff S. Hydroxyl radical production and autoxidative glycosylation. Glucose autoxidation as the cause of protein damage in the experimental glycation model of diabetes mellitus and ageing. Biochem J. 1988;256:205–12.

12. Giugliano D, Ceriello A, Paolisso G. Diabetes mellitus, hypertension and cardiovascular disease: which role for oxidative stress?. Metabolism. 1995;44(3):363–8.

13. Kakkar R, Kalra J, Mantha S, Prasad K. Lipid peroxidation and activity of antioxidant enzymes in diabetic rats. Mol Cell Biochem. 1995;151:113–9.

14. Ozdemirler G, Mehmetcik G, Oztezcan S, Toker G, Sivas A, Uysal M. Peroxidation potential and antioxidant activity of serum inpatients with diabetes mellitus and myocardial infraction. Horm Metab Res. 1995;27:194–6.

15. Jaganjac M, Tirosh O, Cohen G, Sasson S, Zarkovic N. Reactive aldehydes—second messengers of free radicals in diabetes mellitus. Free Radic Res. 2013;47:39–48.

16. Tiwari B, Pandey K, Abidi A, Rizvi S. Markers of oxidative stress during diabetes mellitus. J Biomark. 2013:1–8.

17. Kumar P, Clark M. Textbook of Clinical Medicine. London: Saunders; 2002.

18. Malseed RT. Springhouse Nurses Drug Guide. 6 ed. Philadelphia: Lippincott Williams; 2005.

19. Singh S, Loke Y, Furberg C. Thiazolidinediones and heart failure: a teleo-analysis. Diabetes Care. 2007;30(8):2148–53.

20. White F. Streptozotocin. Cancer Chemotherapy Rep. 1963;30:49–53.

21. Schein P, Cooney D, Vernon M. The use of nicotinamide to modify the toxicity of streptozotocin diabetes without loss of antitumor activity. Cancer Res. 1967;27:2324–32

22. Schein PO, Connel M, Blom J. Clinical antitumor activity and toxicity of streptotocin. Cancer. 1974;34:993–1000.

23. Dorr RT, Fritz WL. Cancer chemotherapy Hand book. London: Kinapton; 1980. 632–37 p.

24. Bonner-Weir S, Trent D, Honey R, Weir G. Responses of neonatal rat islets to streptozotocin: limited B-cell regeneration and hyperglycemia. Diabetes. 1981;30(1):64–9.

25. Hayashi K, Kojima R, Ito M. Strain differences in the diabetogenic activity of streptozotocin in mice. Biol Pharm Bull. 2006;29(6):1110–9.

26. Fadillioglu E, Kurcer Z, Parlakpinar H, Iraz M, Gursul C. Melatonin treatment against remote open injury induced by renal ischemia reperfusion injury in diabetes mellitus. Arch Pharm Res. 2008;31(6):705–12.

27. Etuk E. Animal models for studying diabetes mellitus. Agric Biol J N Am 2010;1(2): 130–4.

28. Sakata N, Yoshimatsu G, Tsuchiya H, Egawa S, Unno M. Animal models of diabetes mellitus for islet transplantation. Exp Diabetes Res. 2012.

29. Rakieten N, Rakieten M, Nadkarni M. Studies on the diabetogenic action of streptozotocin (NSC-37917) Cancer Chemother Rep. 1963;29:91–8.

30. Arison R, Ciaccio E, Glitzer M, Cassaro J, Pruss M. Light and electron microscopy of lesions in rats rendered diabetic with streptozotocin. Diabetes. 1967;16:51–6.

31. Lenzen S, Tiedge M, Jorns A, Munday R. Alloxan derivatives as a tool for the elucidation of the mechanism of the diabetogenic action of alloxan. Lessons from animal diabetes. 1996:113–22.

32. Tjälve H, Wilander E, Johansson E. Distribution of labelled streptozotocin in mice: uptake and retention in pancreatic islets. J Endocrinol. 1976;69:455–6.

33. Karunanayake E, Baker J, Christian R, Hearse D, Mellows G. Autoradiographic study of the distribution and cellular uptake of (14C)-streptozotocin in the rat. Diabetologia. 1976;12:123–8.

34. Ledoux S, Wilson G. Effects of streptozotocin on a clonal isolate of rat insulinoma cells. Biochim Biophys Acta. 1984;804:387–92

35. Schnedl W, Ferber S, Johnson J, Newgard C. STZ transport and cytotoxicity. Specific enhancement in GLUT2-expressing cells. Diabetes. 1994;43:1326–33.

36. Rerup C. Drugs producing diabetes through damage of the insulin secreting cells. Pharmacol Rev. 1970;22:485–518

37. Weiss R. Streptozocin: a review of its pharmacology,efficacy, and toxicity. Cancer Treat Rep. 1982;66:427–38.

38. Ledoux S, Woodley S, Patton N, Wilson G. Mechanisms of nitrosourea-induced beta-cell damage. Alterations in DNA. Diabetes 1986;35:866–72.

39. Murata M, Takahashi A, Saito I, Kawanishi S. Sitespecific DNA methylation and apoptosis: induction by diabetogenic streptozotocin. Biochem Pharmacol. 1999;57:881–7.

40. Konrad R, Kudlow J. The role of O-linked protein glycosylation in beta-cell dysfunction. Int J Mol Med. 2002;10:535–9.

41. Yamamoto H, Uchigata Y, Okamoto H. Streptozotocin and alloxan induce DNA strand breaks and poly (ADP-ribose) synthetase in pancreatic islets. Nature. 1981;294:284–6.

42. Uchigata Y, Yamamoto H, Kawamura A, Okamoto H. Protection by superoxide dismutase, catalase, and poly (ADPribose) synthetase inhibitors against alloxan- and streptozotocininduced islet DNA strand breaks and against the inhibition of proinsulin synthesis. J Biol Chem. 1982;257:6084–8.

43. Sandler S, Swenne I. Streptozotocin, but not alloxan,induces DNA repair synthesis in mouse pancreatic islets in vitro. Diabetologia. 1983;25:444–7.

44. Bennett R, Pegg A. Alkylation of DNA in rat tissues following administration of streptozotocin. Cancer Res. 1981;41:2786–90.

45. Wilson G, Hartig P, Patton N, LeDoux S. Mechanisms of nitrosourea-induced beta-cell damage. Activation of poly (ADP-ribose) synthetase and cellular distribution. Diabetes. 1988;37:213–6.

46. Zafar M, Naqvi SN-u-H, Ahmed M, Khani ZAK. Altered liver morphology and enzymes in streptozotocininduced diabetic rats. Int J Morphol. 2009;27(3):719–25.

47. Zafar M, Naqvi SN-u-H, Ahmed M, Khani ZAK. Altered kidney morphology and enzymes in streptozotocininduced diabetic rats. Int J Morphol. 2009;27(3):783–90.

48. Akpan HD, Ekaidem I. Modulation of Immunological and Hematological Disturbances of Diabetes Mellitus by Diets Containing Combined Leaves of Vernonia amygdalina and Gongronema latifolium. British Journal of Applied Science & Technology. 2015;6(5):534–44.

49. Keskin E, DÖNMEZ N, Kılıçarslan G, Kandir S. Beneficial Effect of Quercetin on Some Haematological Parameters in Streptozotocin-Induced Diabetic Rats. Bulletin of Environment, Pharmacology and Life Sciences. 2016;5(6):65–8.

50. Peelman F, Waelput W, Iserentant H, Laven D, Eyckerman S, Zabeau L, et al. Leptin: linking adipocyte metabolism with cardiovascular and autoimmune diseases. Progress in Lipid Research. 2004;43:283–301.

51. Pertynska-Marczewska M, Kiriakidis S, Wait R, Beech J, Feldmann M, Paleolog E. Advanced glycation end products upregulate angiogenic and pro-inflammatory cytokine production in human monocyte/macrophages. Cytokine. 2004;28:35–47.

52. Shurtz-Swirski R, Sela S, Herskovits AT, Shasha SM, Shapiro G, Nasser L, et al. Involvement of peripheral polymorphonuclear leukocytes in oxidative stress and inflammation in type 2 diabetic patients. Diabetes Care. 2004;24:104–10.

53. Zafar M, Naqvi S. Effects of STZ-induced diabetes on the relative weights of kidney, liver and pancreas in albino rats: a comparative study. Int J Morphol. 2010;28:135–42.

54. Ahmed O, Mahmoud A, Abdel-Moneim A, Ashour M. Antidiabetic effects of hesperidin and naringin in type 2 diabetic rats. Diabetol Croat. 2012;41:53–67.

55. Kim J, Choi J, Lee M, Kang K, Paik M, Jo S, et al. Immunomodulatory and antidiabetic effects of a new herbal preparation (HemoHIM) on streptozotocin-induced diabetic mice. Evid Based Complementary Alternat Med. 2014:1–8.

56. Salih ND, Kumar GH, Noah RM, Muslih RK. The effect of streptozotocin induced diabetes mellitus on liver activity in mice Global Journal on Advances in Pure & Applied Sciences 2014;3:67–75.

57. Nagarchi K, Ahmed S, Sabus A, Saheb SH. Effect of Streptozotocin on Glucose levels in Albino Wister Rats J Pharm Sci & Res 2015;7(2):67–9.

58. Dacie J, Lewis S. Practical haematology. 7 ed: Edinburgh:ELBS with Churchill Livingstone; 1991.

59. Alexander R, Grifiths J. Basic biochemical methods. 2 ed. New York: Wiley-Liss; 1993.

60. Osim E, Akpogomeh B, Ibu J, Eno A. Experimental physiology manual. 3 ed: Calabar:University of Calabar,Dept Physiol; 2004.

61. Mcanuff M, F. Omoruyia, Morrison E, Asemote H. Hepatic function enzymes and lipid peroxidation in STZ-induced diabetic rats fed bitter yam (Dioscorea polygonoides) steroidal sapogenin extract. Diabetol Croat. 2003;32:17–23.

62. Kennedy L, Baynes J. Non-enzymatic glycosylation and the chronic complications of diabetes: An Overview.

63. Low A, Nickander K, Tritschler J. The role of oxidative stress and antioxidant treatment in experimental diabetic neuropathy. Diabetes. 1977;46:38–41.

64. Arora S, Ojha S, Vohora D. Characterisation of streptozotocin induced diabetes mellitus in swiss albino mice. Global J Pharmacol. 2009;3:81–4.

65. Kolb H, Kroneke D. IDDM: lessons from the low dose streptozotocin model in mice. Diabetes Rev. 1993;1:116–26.

66. Kang N, Alexander G, Park JK, Maasch C, Buchwalow I, Luft LC, et al. Differential expression of protein kinase C isoforms in streptozotocin-induced diabetic rats. Kidney Int. 1999;56(5):1737–50.

67. Akbarzadeh A, Norouzian D, Mehrabi MR, Jamshidi S, Farhangi A, Verdi AA, et al. Induction of diabetes by Streptozotocin in rats. Indian Journal of Clinical Biochemistry. 2007;22(2):60–4.

68. Koga T, Keiko M, Terao J. Protective effet of vitamin E analog, phosphatidylchromanol, against oxidative hemolysis of human erythrocytes. Lipids. 1980;33.

69. Sen S, Kar M, Roy A, Chakraborti A. Effect of nonenzymatic glycation on function and structural properties of hemoglobin. Biophys Chem. 2005;113:289–98.

70. Kolanjiappan K, Manoharan S, Kayalvizhi M. Measurement of erythrocyte lipids, lipid peroxidation, antioxidants and osmotic fragility in cervical cancer patients. Clin Chim Acta 2002;326:143–9.

71. Weiss G, Goodnough L. Anemia of chronic disease. N Engl J Med. 2005;352:1011–23.

72. Akindele O, Babatunde A, Chinedu F, Samuel O, Oluwasola C, Oluseyi A. Rat model of food induced non-obese-type 2 diabetes mellitus; comparative pathophysology and histopathology. International Journal of Physiology, Pathophysiology and Pharmacology. 2012;4(1):51–8.

73. Stookey JD, Burg M, Sellmeyer DE, Greenleaf JE, Arieff A, Hove LV, et al. A proposed method for assessing plasma hypertonicity in vivo. European Journal of Clinical Nutrition. 2007;61:143–6.

74. Domenico ID, Ward DM, Kaplan J. Regulation of iron acquisition and storage:consequences for iron-linked disorders. Nat Rev Mol Cell Biol. 2008;9:72–81.

75. Hotamisligil G. Inflammation and metabolic disorders. Nature 2006;444:860–7.

76. Korpinen E, Groop P, Fagerudd J, Teppo A, Akerblom H, Vaarala O. Increased secretion of TGF-beta1 by peripheral blood mononuclear cells from patients with type 1 diabetes mellitus with diabetic nephropathy. Diabet Med. 2001;18:121–5.

77. Kedziora-Kornatowska K. Production of superoxide and nitric oxide by granulocytes in non-insulin-dependent diabetic patients with and without diabetic nephropathy. IUBMB Life. 1999;48:359–62.

78. Shanmugam N, Reddy M, Guha M, Natarajan R. High glucose-induced expression of proinflammatory cytokine and chemokine genes in monocytic cells. Diabetes. 2003;52:1256–64.

79. Frühbeck G. Intracellular signalling pathways activated by leptin. Biochem J. 2006;393:7–20.

80. Baumann H, Morella KK, White DW, Dembski M, Bailon PS, Kim H, et al. The full-length leptin receptor has signaling capabilities of interleukin 6-type cytokine receptors. Proc Natl Acad Sci USA. 1996;93:8374–8.

81. Kozlov I, Novitski V, Baìkov A. Kinetics of blood leukocytes in mice with alloxan diabetes. Biull Eksp Biol Med. 1995;120:33–5.

82. Latry P, Bioulac-Sage P, Echinard E, Gin H, Boussarie L, Grimaud J, et al. Perisinusoidal fibrosis and basement membrane-like material in the livers of diabetic patients. Hum Pathol. 1987;18(8):775–80.

83. Das A, Padayatti P, Paulose C. Effect of leaf extract of Aegle marmelose (L.) Correa ex Roxb. on histological and ultrastructural changes in tissues of streptozotocin induced diabetic rats. Indian J Exp Biol. 1996;34(4):341–5.

84. Harrison S. Liver disease in patients with diabetes mellitus. J Clin Gastroenterol. 2006;40(1):68–76.

85. Houstis N, Rosen E, Lander E. Reactive oxygen species have a causal role in multiple forms of insulin resistance. Nature. 2006;440(7086):944–8.

86. Poitout V, Robertson R. Glucolipotoxicity: fuel excess and beta-cell dysfunction. Endocr Rev. 2008;29(3):351–66.

87. West I. Radicals and oxidative stress in diabetes. Diabet Med. 2000;17(3):171–80.

88. Ozcan U, Cao Q, Yilmaz E, Lee A, Iwakoshi N, Ozdelen E. Endoplasmic reticulum stress links obesity, insulin action, and type 2 diabetes. Science. 2004;306:457–61.

89. Hatcher H, Planalp R, Cho J, Torti S. Curcumin from ancient medicine to current clinical trials. Cell Mol Life Sci. 2008;65:1631–52.

90. Kushner I. The phenomenon of the acute phase response. Ann NY Acad Sci. 1989;557:39–46.

91. Schultz D, Arnold P. Properties of four acute phase proteins: C-reactive protein, serum amyloid aprotein,α1-acid glycoprotein, and fibrinogen. Seminars in Arthritis and Rheumatism. 1990;20(3):129–47.

92. Yuki M, Machida N, Sawano T, Itoh H. Investigation of serum concentrations and immunohistochemical localization of α1-acid glycoprotein in tumor dogs. Veterinary research communications. 2011;35(1):1–11.

93. Kyle R, Katzmann J, Lust J, Dispenzieri A. Clinical indications and applications of electrophoresis and immunofixation. In Manual of Clinical Laboratory Immunology. 6 ed. Washington DC: ASM Press; 2002.

94. Jacobs SD, DeMott RW, Oxley KD. Laboratory test handbook. 3 ed: Lexi comp; 2004.

95. Pagana K, Pagana T. Mosby’s manual of diagnostic and laboratory tests. Elsevier Health Sciences. 2013.

